# Active and poised promoter states drive folding of the extended *HoxB* locus in mouse embryonic stem cells

**DOI:** 10.1101/111773

**Authors:** Mariano Barbieri, Sheila Q. Xie, Elena Torlai Triglia, Inês de Santiago, Miguel R. Branco, David Rueda, Mario Nicodemi, Ana Pombo

## Abstract

Gene expression states influence the three-dimensional conformation of the genome through poorly understood mechanisms. Here, we investigate the conformation of the murine *HoxB* locus, a gene-dense genomic region containing closely spaced genes with distinct activation states in mouse embryonic stem (ES) cells. To predict possible folding scenarios, we performed computer simulations of polymer models informed with different chromatin occupancy features, which define promoter activation states or CTCF binding sites. Single cell imaging of the locus folding was performed to test model predictions. While CTCF occupancy alone fails to predict the in vivo folding at genomic length scale of 10 kb, we found that homotypic interactions between active and Polycomb-repressed promoters co-occurring in the same DNA fibre fully explain the HoxB folding patterns imaged in single cells. We identify state-dependent promoter interactions as major drivers of chromatin folding in gene-dense regions.

---

The link between gene expression states and long-range chromatin contacts between different transcription units is a major open question in our understanding of gene regulation (**Fig. 1a**). Imaging approaches suggest that active transcription units cluster in sites of transcription, called transcription factories^1,2^, whereas Polycomb-repressed genes are found to associate with Polycomb bodies or poised transcription factories^3-6^. Genes in the same metabolic pathways can co-localise with each other, especially in specialized cell types, although at low frequency across cell populations^7,8^. Although the molecular mechanisms that underlie promoter co-associations and their functional purpose remain unclear, they suggest that gene activation states may help to establish cell-type specific chromatin folding configurations that partition active from Polycomb-repressed chromatin domains. CTCF binding has important roles in the formation of chromatin loops and enhancer-promoter interactions^9-14^, and has been proposed to organise chromatin domains through loop extrusion mechanisms in gene-poor areas^15^ and to help insulate spreading of active marks into Polycomb repressed domains^16^. However, CTCF contribution to the large scale folding of more complex, gene-dense regions remains unknown in particular regions populated with active and Polycomb-repressed loci.

**Figure 1.**
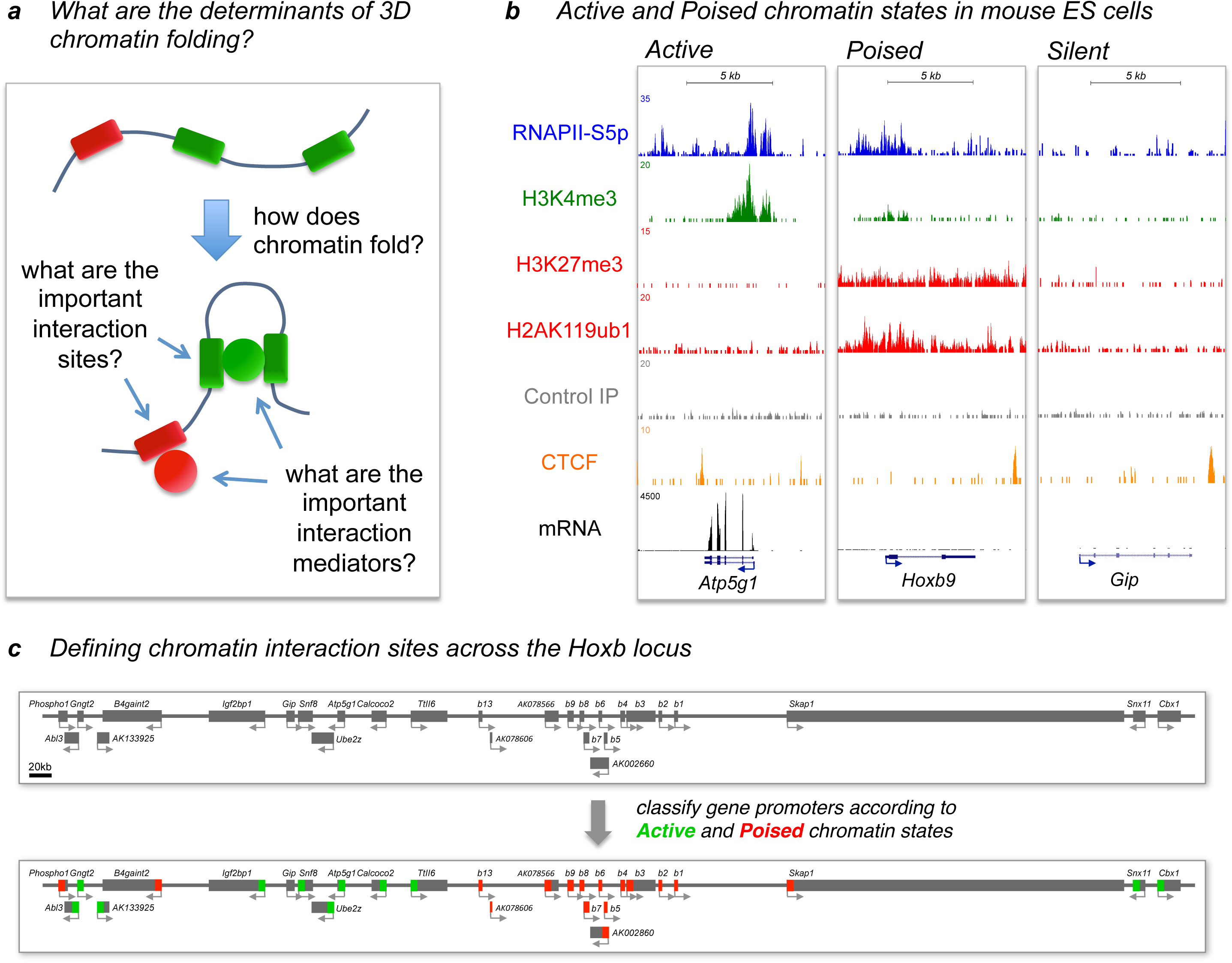
Classification of gene promoter states across the *HoxB* locus in mouse ES cells. **a)** Our study focuses on understanding the molecular mechanisms that underlie chromatin folding and the relationship with gene activation states. **b)** Gene activation states can be identified using chromatin occupancy maps produced by ChIP-seq for RNAPII-S5p, H3K4me3, H3K27me3 and H2AK119ub1 in mouse ES cells. Control (mock IP), CTCF binding and mRNA-seq profiles are also shown. Examples are shown for active gene *Atp5g1,* Polycomb-repressed gene *Hoxb9* and silent gene *Gip,* all present within the *HoxB* cluster. **c)** Classification of gene promoter states across the extended *HoxB* locus according to RNAPII occupancy and Polycomb repressive marks as Active, Poised and Silent states.

Mouse ES cells lack lamin A and have lower levels of heterochromatin than differentiated cells^17^. Although some reports suggest that ES cell chromatin is more dynamic than the chromatin of differentiated cells, with higher exchange of linker histone H1 and nucleosome dynamics^18^, other studies find similar kinetics^19,20,21^. Nevertheless, chromosomes retain their territorial conformation in ES cells, and RNA polymerase II (RNAPII) organizes itself in active and poised transcription factories ^3,4,22,23^. Active transcription compartments, or transcription factories^1,8,24–26^, associate with active genes and are characterized by the presence of RNAPII complexes phosphorylated on both Serine5 and Serine2 residues of C-terminal domain of the largest subunit, Rpb1^2–4^ Poised transcription factories contain Polycomb and RNAPII phosphorylated on Serine5 only, and co-localise with Polycomb-repressed genes^3,4,27^. Polycomb repression plays major roles in pluripotent cells through the silencing of early developmental genes, which are kept in a poised state ready for induction in response to signals that commit cells into specific embryonic lineages^5,28,29^. Polycomb complexes modify histone tails by di-or trimethylation of histone H3K27 residues and monoubiquitinylation of histone H2AK119 residues. Polycomb-repressor complexes can induce chromatin folding independently of catalytic activity^30-32^. The co-associations of active genes or Polycomb-repressed genes have so far been studied separately, without assessing whether one type of association might have predominant contributions to chromatin folding at specific loci.

To further understand the underlying mechanisms of chromatin folding and identify candidate drivers, we focus our study on a 1Mb gene-dense region of the murine genome, centred on the *HoxB* locus, the least studied of the *Hox* loci at the level of chromatin topology. The *HoxB* locus is a complex locus containing closely spaced genes, in two states of transcriptional activation in mouse ES cells, active or Polycomb-repressed^3^.

To explore the mechanisms of chromatin folding in the *HoxB* locus and to dissect different possible folding scenarios, we use a polymer physics approach, the Strings &Binders Switch (SBS)^33-35^ model. We considered different scenarios of chromatin contacts driven by: *(i)* different promoter states (Active or Polycomb-repressed); *(ii)* only driven by the presence of RNAPII-S5p/H3K4me3 (without considering the contribution of Polycomb); or *(iii)* CTCF binding factor occupancy. To discriminate different folding scenarios of the *HoxB* locus, we performed single cell imaging using fluorescence *in situ* hybridization (FISH) in mouse ES cells. We find that the active and Polycomb-repressed promoter states play separate but simultaneous roles in the folding of the *HoxB* locus at the genomic length scale of tens of kilobases, whereas CTCF occupancy alone does not explain folding at this scale.

## Results

### Identifying chromatin states across the HoxB locus in mouse ES cells

We considered a 1Mb genomic region centred on the *HoxB* gene cluster (chr11: 95685813-96650449), containing *Hoxb1-b9, b13* and a total of 28 RefSeq genes (**Fig. 1b,c**). To define the genomic position of CTCF binding sites and gene promoter states within such region, we mined published datasets of chromatin occupancy or published classifications ^27,36,37^ (**Supplementary Table 1**).

The Active promoter state was defined by the presence of RNAPII-S5p and H3K4me3 within a 2kb window centred on their transcription start site (TSS), and by the absence of Polycomb-repressive marks (e.g. *Atp5g1* gene; **Fig. 1b,c**), as described previously^3^. The Polycomb-repressed (poised) promoter state was defined by the presence of RNAPII-S5p, H3K4me3, and the presence of at least one of the Polycomb-repressive marks H2AK119ub1 or H3K27me3 at TSS regions, but not mRNA expression (e.g. *Hoxb9*; **Fig. 1b,c**). We find that 27 promoters can be classified into either Active or Poised states, while the promoter of the *Gip* gene was not found occupied by RNAPII-S5p, H3K4me3 or H3K27me3, and was classified as inactive (complete classification in **Supplementary Table 2**). As an additional folding scenario, we considered a possible contribution of open promoter states, marked by RNAPII-S5p and H3K4me3 occupancy, irrespectively of their Polycomb state (**Supplementary Fig. 1a-c**). Finally, we used a published list of CTCF binding sites, which are abundant in the full 1Mb length of the locus, as candidate drivers for the folding of the extended *HoxB* locus (**Supplementary Fig. 1d**).

### Modelling different folding conformations of the *HoxB* locus

To dissect the folding mechanisms of the *HoxB* locus, we used the SBS polymer model^33,35^ to simulate the effects of the different possible molecular determinants. The SBS model simulates a scenario where chromatin conformation is shaped by chromatin interactions at specific genomic locations (or binding sites; **Fig. 2a**, red and green) with cognate binding molecular factors. The SBS model has been previously used to successfully describe Hi-C data^33^, the relationship between genomic distance and the frequency of chromatin contacts measured by FISH in mammalian and fly nuclei^33,38^, co-localisation events of *Xist* loci during X inactivation in female mouse cells^39,40^, and more^41,42^.

**Figure 2.**
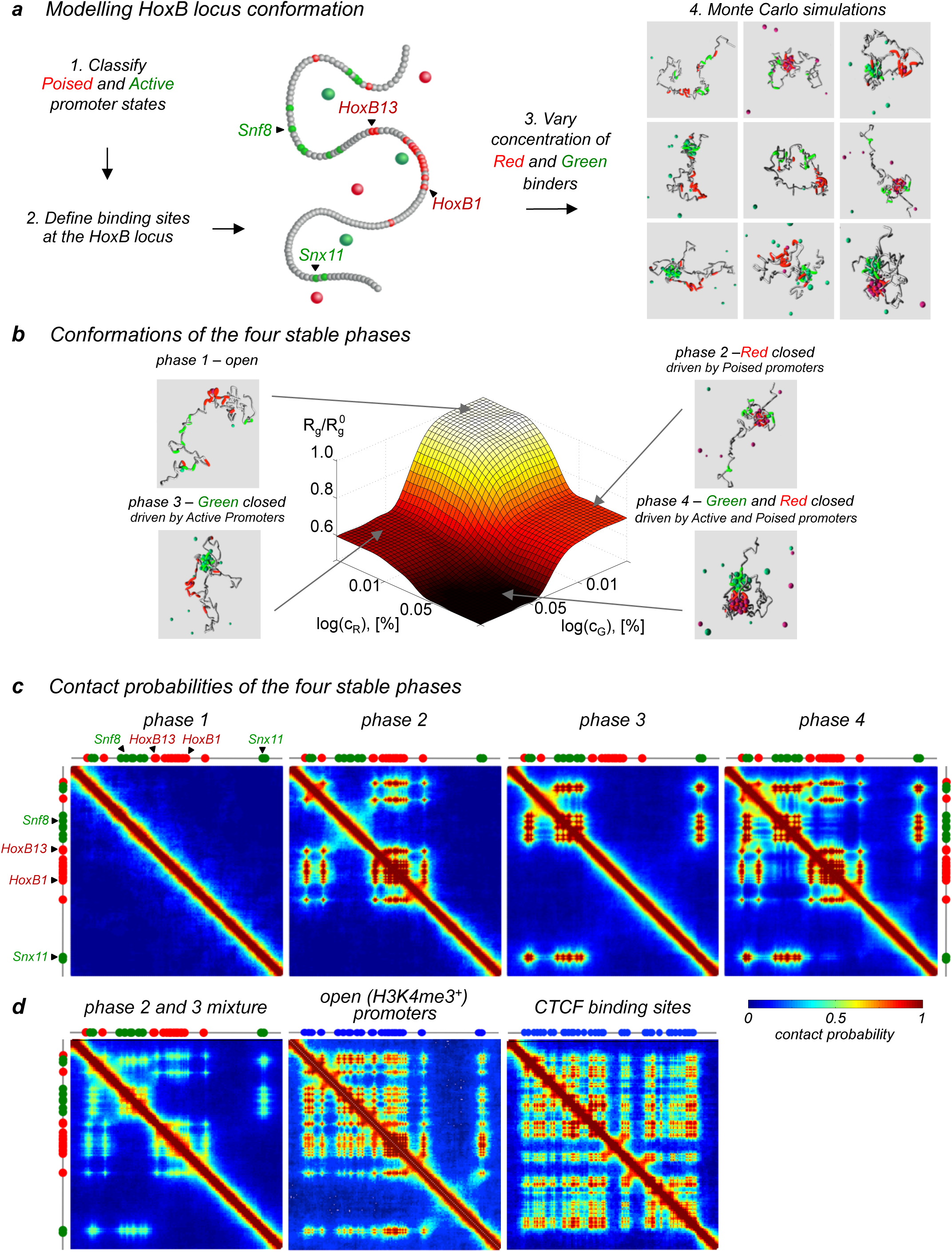
The SBS polymer model predicts folding of the *HoxB* locus under different biological scenarios. **a)** The 1Mb extended *HoxB* locus is mapped at a 7.6kb resolution into an SBS bead-chain polymer model, informed with binding sites that represent gene promoter states classified as active (green), poised (red) or silent (grey). Active and poised binding sites can bind ‘active’ (green) and ‘poised’ (red) binders, respectively. Monte Carlo simulations from polymer physics identify the equilibrium conformations of locus folding. **b)** The system phase diagram identifies the stable architectural classes of the model, which correspond to its thermodynamics phases, defined by concentration of binders. **c)** Different contact probability patterns are identified in the four thermodynamic phases. **d)** Additional contact probability patterns from control models: a 50%-50% mixed population of phase 2 and phase 3 models; folding driven by open promoters, considering exclusively the presence of RNAPII-S5p and H3K4me3; and, folding driven by CTCF-dependent contacts.

The 1Mb long genomic region containing the *HoxB* locus was modelled within the SBS, as a self-avoiding polymer bead chain^33,35^ (see Online Methods). To achieve reasonable computer simulation times, we defined a polymer with a total of 152 beads, where each polymer bead corresponds to 7.6 kb; 12 inert beads were added on each polymer tail to counterbalance finite size effects. The genomic length of each bead was chosen to avoid the presence of genes with different promoter states within the same bead. For computational efficiency, we simulate the polymer on a square lattice with periodic boundary conditions. The locations and types of binding sites and binding factors are chosen from the chromatin states described above.

We first considered polymers where the potential binding sites are defined by whether they contain Active and/or Poised promoters (**Fig. 1c**), and where contacts can only be mediated through the co-associations of beads with affinity to the same kind of binding site. We classified binding sites according to promoter states only (and not whole coding regions), because promoters are the gene region that are most abundantly associated with RNA polymerase II and Polycomb repressor complexes^43^. In such models, polymer beads containing the coordinates of gene promoters reflect the state of activation of genes across the locus (**Fig. 2a**), namely Active (green) state of activation and Poised (or Polycomb-repressed; red) state of activation. All the other polymer beads are in the inert (grey) state, and they do not interact except for excluded volume effects. Specific binding is promoted by two different kinds of Brownian molecules: ‘active’ and ‘poised’ binding factors (see **Fig. 2a**, green and red free beads, respectively). Each binding factor is only allowed to bind to binding sites of the same colour, and can bind multiple binding sites, up to a maximum of six (the coordination number on the cubic lattice), in the same order of magnitude of the number of polymerases that is suggested to co-exist in a transcription factory^2,22^. To investigate the effects of binding valency of the polymer binding sites, we discuss first the case where interacting polymer beads can be bound simultaneously by up to 6 binders; as a comparison, we later repeated simulations where binding sites can only be bound by a single binder while allowing each binder to bind 6 binding sites.

Using computer simulations, we explored the equilibrium conformational states of such model system. We varied the concentrations of ‘poised’ and ‘active’ binding molecules, *c_R_* and *c_G_*, in a broad, biologically relevant range, and derived the system phase diagram for *E_R_=E_G_=4k_B_T,* a value in the range of real transcription factor binding energies^44,45^ (**Fig. 2b**; see Online Methods). Here, *k_B_* is the Boltzmann constant and *T* the temperature in Kelvin (see Online Methods). The phase diagram obtained, at fixed binder concentrations, varying the energy of binding of polymer beads for ‘red’ and ‘green’ binders, *E_R_* and *E_G_,* has a similar structure (see **Supplementary Fig. 2**). It is well established in polymer physics that the system thermodynamics phases do not depend on specific choices of model details (e.g. polymer length or lattice coordination number) and are, in this sense, fully general^46^. Whereas additional complexities could be considered, in particular to explain chromatin folding at lower genomic resolutions, our present aim was focused on testing to what extent could a simplified model of chromatin folding based only on the biological functionality of gene promoters be sufficient to derive folding scenarios that could be tested experimentally to help understand larger-scale folding mechanisms of a complex locus.

### The model conformational classes and their contact matrices

Polymer modelling of the *HoxB* locus in different concentrations or energies of binding identifies four distinct thermodynamics phases and four corresponding conformational classes (**Fig. 2b** and **Supplementary Fig. 2;** see also Barbieri et al. 2012^33^). In phase 1, polymers are fully open and have random folded conformations, as expected for self-avoiding walk (SAW) polymers. In phase 2, only the red (Polycomb-like) beads fold spontaneously to form a ‘red’ compact globule, while the green polymer binding sites (i.e. Active promoters) float independently in a SAW state. In these conditions, only Polycomb-repressed promoters drive the locus folding. In phase 3, the red beads are open and the green beads form a globule, corresponding to an Active model where only active promoters drive folding. Finally, in phase 4, both red and green beads compete for binding to their specific binding sites and form two separate compact structures with the intervening grey polymer beads looping out.

To help visualize the contact patterns and the frequency across the ensemble of polymers modelled in each phase, we produced contact matrices that represent the pair-wise contact probability of polymer sites (i.e., the probability that they are distant by no more than three lattice units, d_0_). Each thermodynamic phase gives rise to distinct contact matrices (**Fig. 2c**). In phase 1, the fully open phase, the contact matrix has only a diagonal signature, as only neighbouring sites have higher chances of random contacts. Conversely, strikingly different contact patterns emerge in the other three phases, which reflect only red (phase 2), only green (phase 3) or a combination of green and red homotypic contacts (phase 4); bystander effects are also observed.

We also considered three additional cases. First, to explore the effect of heterogeneous chromatin contacts across cell populations or different alleles in the same cell, we considered mixtures of polymers folded in the green-only and red-only states (50%-50% mixture of phases 2 and 3; **Fig. 2d**). We also implemented polymer models representing the other scenarios for *HoxB* folding: a model where polymer binding sites are only linked to ‘open’ promoter states associated with the presence of RNAPII and H3K4me3 (irrespectively of presence of Polycomb), and a model where folding is driven by CTCF binding site locations (**Supplementary Fig. 1c,d**). In the two latter cases, there is only one type of binding factor such that the system phase diagram has only two phases, as expected: a SAW open state and a state where the interacting beads form a globule, compact conformation^47^. The seven contact matrices obtained with the different models have clearly distinct patterns of folding (**Fig. 2c,d**), which can be tested experimentally by visualising the positioning of the candidate interacting regions in mouse ES cells.

### Testing the model predictions

To explore the real architecture of the *HoxB* locus in mouse ES cells, we set out to determine the inter-chromatin distances across the *HoxB* locus in single cells using cryoFISH, a high-resolution imaging approach, which combines FISH with optimal preservation of cellular architecture^48^, followed by confocal microscopy^49^. This approach was previously used to study gene associations with active or poised transcription factories^1,2,4^, to study chromosome volume, intermingling and their correlation with chromosome translocations^48^, and to validate 4C-seq chromatin contacts^50^. We selected fosmid probes that cover ~30 to 40kb genomic regions (corresponding to ~4 beads in the polymer), as these provide high spatial resolution to study the general properties of folding of a 1Mb region and they are detected with optimal sensitivity by cryoFISH^4^.

To identify the minimal number of genomic regions that would be required to discriminate the seven different folding models of the extended *HoxB* locus, we identified five evenly-spaced genomic regions, each covering ~30kb, which have different contact profiles (Fig. 3a, **Supplementary Table 3**). For example, in the models that consider Active and Poised promoters, the two regions containing active (green) genes, *Snf8/Ube2z* and *Snx11/Cbx1*, would contact each other with high probability in phases 3 and 4, but not in phases 1 or 2 (**Supplementary Fig. 3a**). The regions containing poised (red) genes, *Hoxb1/Hoxb2* and *Hoxb13/AK078606,* would contact only in phases 2 and 4. The same set of probes can also distinguish whether green-green and red-red contacts are simultaneous or separate (i.e. phase 4 from a mixture of phases 2 and 3), as in the latter case we would observe similar patterns of contacts, but with lower intensity, and a model based only on promoter occupancy by RNAPII-S5p (**Supplementary Fig. 3b**). Finally, polymer folding driven by CTCF-binding sites alone would result in contacts between *Snf8/Ube2z* (active) and *Hoxb13/AK078606*, and to a lesser extent *Hoxb1/Hoxb2* and Control regions, but not with the distant pair of active genes, *Snx11/Cbx1*. The intervening, Control region lies within the *Skap1* coding region which contains CTCF binding sites but is devoid of Polycomb marks and H3K4me3 (**Supplementary Fig. 4**).

**Figure 3.**
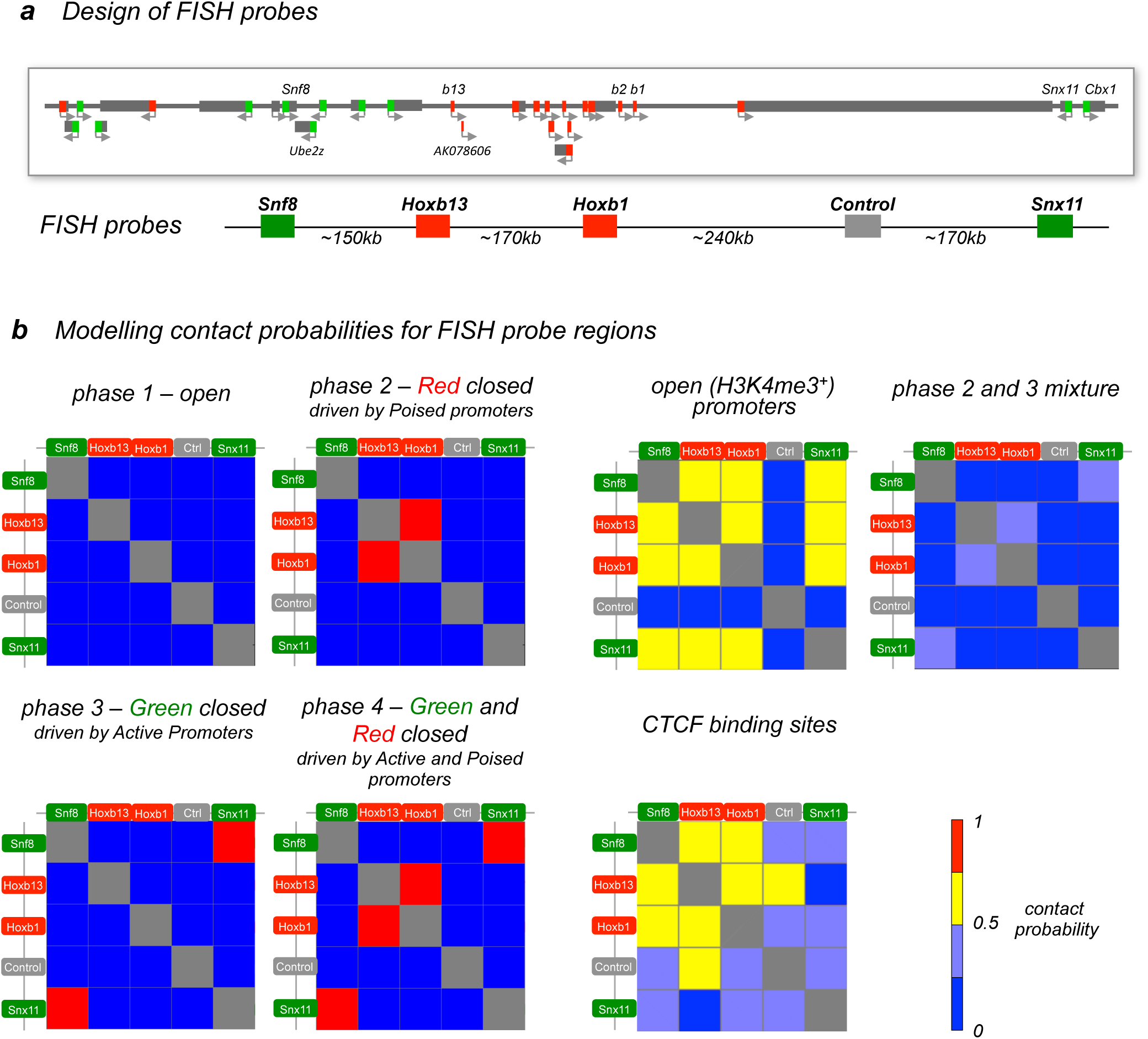
Genomic regions chosen for testing model predictions of *HoxB* locus folding by FISH in ES cells. **a)** Scheme of the positions of the selected FISH probes covering ~35kb regions centred on promoters of *Snf8, Hoxbl, Hoxb13* and *Snx11* genes and at a control region. *Hoxbl* and *Hoxb13* are classified as poised genes (red), whereas Snf8 and Snx11 as active (green). **b)** Mini-contact matrices obtained from modelled polymers of the five genomic regions chosen for experimental testing have clearly different contact maps that correspond to very distinct mechanisms of folding.

To allow direct comparisons between distances derived from cryoFISH experiments (performed on ~150nm thick cryosections cut in random orientations) and the 3D polymer models generated in the different conditions, we simulated the sectioning process on the ensemble of polymer models by cutting random virtual ‘slices’ through our simulated polymers with thickness equivalent to the cryoFISH experiments (see Online Methods). To simulate the imaging of cryosections and distance measurement in the plane of the sections, we determined the inter-locus distances between the centres of mass of each genomic region in the polymer models, considering the projection of the 3D distance on the plane of the section. As expected, all the mini-matrices generated from polymers modelled according to the different scenarios make different predictions of folding of the *HoxB* locus (**Fig. 3b**).

### Measuring chromatin contacts within the *HoxB* locus in mouse ES cells by cryoFISH

To determine the physical distances in the nucleus of mouse ES cells between the five *HoxB* locus regions selected based on the polymer modelling, we performed cryoFISH for all ten pairwise combinations of the five probes. The five regions are separated by only 150–730 kb (**Fig. 4a**), making high-resolution cryoFISH particularly relevant for these analyses, as it uses thin cryosections (~150 nm) to improve the z axis diffraction limit. Visual inspection of cryoFISH images (**Fig. 4a**) suggested higher levels of homotypic co-localisation between the two probes marking the two Poised genomic regions (*Hoxb1/Hoxb2* and *Hoxb13/AK078606*) and the two probes marking the two Active genomic regions (*Snf8/Ube2z* and *Snx11/Cbx1*), than between other probe combinations.

**Figure 4.**
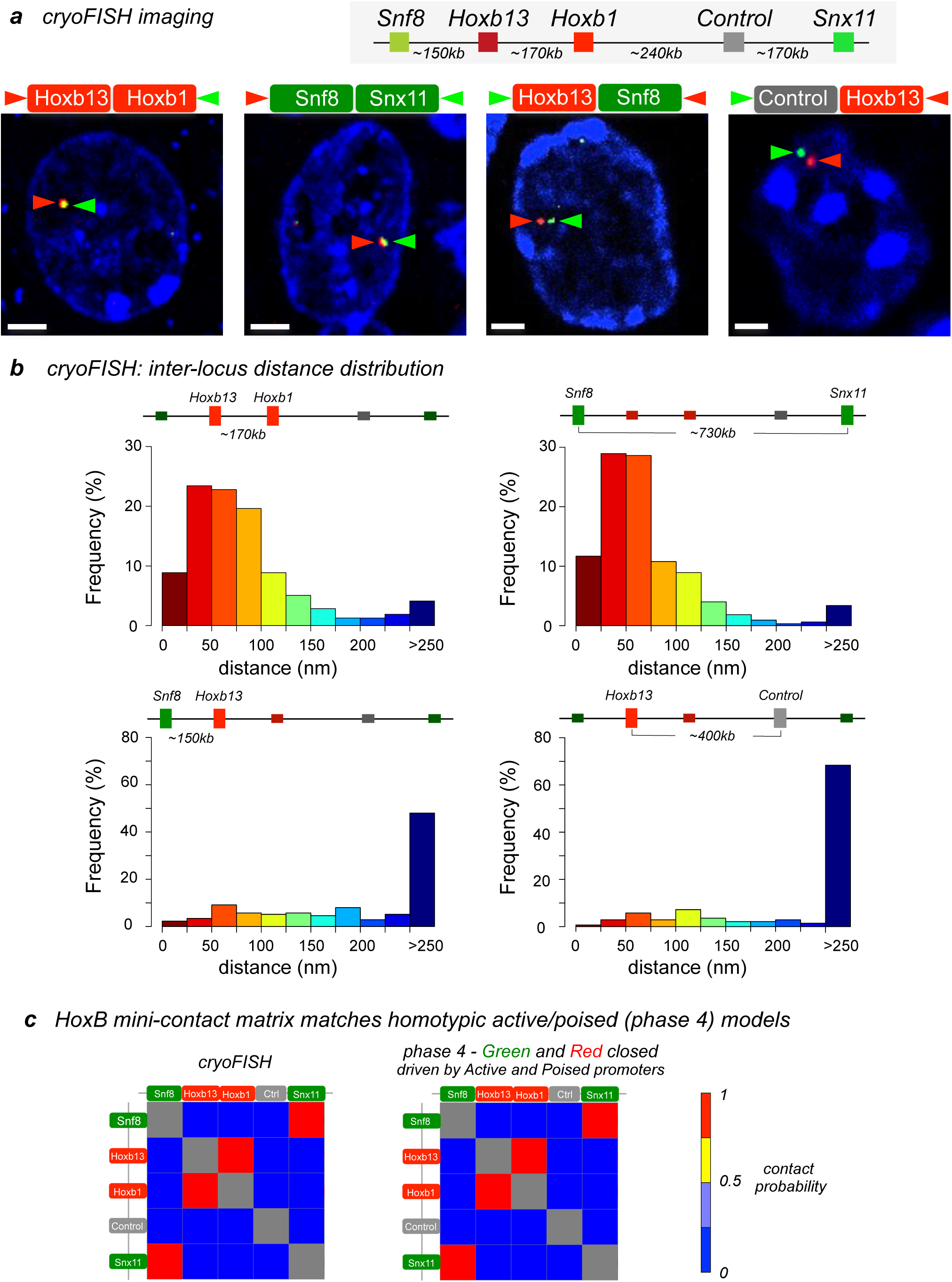
Imaging the extended *HoxB* locus by FISH identifies a folding pattern dependent on homotypic contacts between active and poised promoters, as predicted by the SBS model. **a)** Representative cryoFISH images of *HoxB* locus folding. Top, schematic of the genomic positioning of the cryoFISH probes over *HoxB* locus. Cryosections (150 nm thick) were hybridized with probes specific to *Hoxb1, Hoxb13, Snx11, Snf8* and control. Coloured boxes around gene names represent Active (green) and Poised (red) promoter states. Coloured arrowheads pointing at gene names indicate the pseudo-colour chosen to represent each probe signal in the image. Nucleic acids were counterstained with TOTO-3 (blue). Bars, 2 μm. **b)** Histograms of inter-locus physical distances between different pairs of probes. The average distances between poised (top left; *Hoxb1, Hoxb13)* or active promoter regions (top right; *Snf8, Snx11)* are significantly shorter (<150 nm) than between other pairs of loci (e.g., bottom left; *Snf8, Hoxb13)* or loci relative to control (bottom right; *Snx11,* control), most often found at >250 nm. **c)** The contact mini-matrix determined experimentally by cryoFISH imaging is equivalent to the polymer phase 4 prediction of the SBS model obtained by slicing polymer ensembles, suggesting that distinct homotypic interactions between active and poised regions of the *HoxB* locus are a major folding feature.

To obtain a quantitative description of the *HoxB* locus folding, we collected a large dataset of cryoFISH images by confocal microscopy, and identified the position of each genomic region within the *HoxB* locus using a semi-automated macro implemented in ImageJ (**Supplementary Fig. 5a;** see Online Methods). For each of the ten probe pairs, we collected images from 1,535-2,587 nuclear profiles (adding to a total of ~20,000 nuclear profiles). As expected due to the thin sectioning and random orientation, most nuclear profiles do not contain the *HoxB* locus (**Supplemetary Fig. 5a**), resulting in a dataset of ~200 inter-locus distances per probe pair (~2000 inter-locus distances for all probe pairs; **Supplementary Fig. 5b**). The probes produced give balanced frequency of locus detection per nuclear slice (on average 14±3%, as expected in ~150 nm sections; **Supplementary Fig. 5b**). Red and green signals were detected with proportional frequency within the same FISH experiment; difference to the average detection was 7±4% most likely due to differences in probe length, their labelling efficiency and potentially locus compaction. The small variability in detection between cryoFISH experiments may result from small differences in section thickness.

We then measured the distances between the centres of mass of each probe signal (see Online Methods). We find that the distributions of distances between genomic regions from the same class (homotypic active-active or poised-poised pairs) are strikingly narrower than distributions of pairs from different classes (heterotypic pairs) or pairs including the control region (**Fig. 4b** and **Supplementary Fig. 6**), as also shown by their mean distances (**Supplementary Fig. 5b**). The centres of mass of signals between the two poised regions (*Hoxb1/Hoxb2* and *Hoxb13/AK078606)* are found within 100 nm in the vast majority (~75%) of alleles, identifying a frequent and spatially close contact between these two regions that persists across the cycling population of ES cells. This observation agrees with reports of stable associations between Polycomb repressed domains^51^. Co-localisation between *Hoxb1* and *Hoxb9* had been previously shown in mouse ES cells by 3D-FISH yielding an average distance of 100 nm between the two genomic regions^52^, compatible with our findings of average ~70 nm. Remarkably, although the two probes covering active genes (*Snf8/Ube2z* and *Snx11/Cbx1*) are separated by 730 kb in linear genomic sequence, their physical distance and frequency of contact are equally close (~80 nm) and frequent (~75%), respectively, indicating that these two active regions co-localise in the vast majority of ES cells in the population, often for both alleles. In contrast, for all the remaining pairs of regions, we find only a small proportion (10-25%) of the distances at <100 nm, with the vast majority (65-90%) separated by >250 nm. These results identify the spatial compartmentalization of Active and Polycomb repressed promoter regions, across hundreds of kilobasepairs within the extended *HoxB* locus, and show the power of cryoFISH in fine mapping genomic distances even between regions closely spaced along the linear genome. We identify critical co-localisation distances below ~100 nm, for sub-regions within the *HoxB* locus, which are promptly distinguishable from distances above 250 nm detected for regions within the same locus that do not co-localise.

### Comparing polymer models with locus distances in mouse ES cell nuclei

To compare the folding of the *HoxB* locus observed in ES cells with the seven model predictions, we produced a cryoFISH contact matrix by calculating the frequency of FISH probe co-localisation (**Fig. 4c**; using co-localisation threshold of 105 nm; see Online Methods). The matrix is directly comparable with the matrices shown in **Fig. 3b** produced by slicing 3D polymer models. Strikingly, we find that the experimental matrices exactly match phase 4 of the model which considers simultaneous contacts between active and between poised gene promoters, where both Active (green) and Polycomb (red) beads can contact their corresponding green and red beads, respectively, within the same polymer. We find that the pattern of folding is the same, and also the frequency of contacts is similar, with ~75% of active and poised regions found within 105 nm. Conversely, the alternative scenarios obtained by polymer modelling are incompatible with the experimental FISH data. In particular, the mixed population scenario where contacts between Active regions and between Poised regions are mutually exclusive in the same polymer and occur in separate alleles or different cells (**Fig. 3b**), is not compatible with the high frequency of chromatin contacts identified experimentally (**Fig. 4c**), irrespectively of an overall identical pattern of contacts between the two cases. Contacts mediated by CTCF binding sites alone (**Fig. 3b**) are particularly unable to explain the matrices of contacts detected experimentally across the gene-dense *HoxB* locus, although they may contribute to the overall folding as seen in the other *Hox* loci^16^, and may have deterministic roles in gene-poor areas, as those recently explored^15^. Our detailed analyses of physical distances across the *HoxB* locus suggest that homotypic (Active-Active and Poised-Poised) contacts are highly predominant within ES cell nuclei, and are not explained by independent contacts in separate cells or even separate alleles within the same cells.

To explore to what extent were the predictions of our simple models able to explain further details of folding, we compared the entire experimental locus distance distributions observed experimentally with the distance distributions predicted for the ensemble of polymers in phase 4 (**Fig. 5** and **Supplementary Fig. 7**). We found that the distance distributions determined in ES cell nuclei are in good agreement with model predictions for phase 4 (**Fig. 5** and **Supplementary Fig. 7**). In contrast, when the mixture model is considered where Active-Active or Poised-Poised interactions occur in separate polymers (50%-50% phase 2 and 3), the distance distributions become bimodal and include much larger separations which were not observed experimentally (**Fig. 5**). The precise details of the distance distributions measured by cryoFISH are less well recapitulated for the distances between heterotypic than the homotypic genomic regions, suggesting that other folding properties or binding sites will need to be considered in future to explain further details of the *HoxB* locus folding. Features that can play a role in fine-tuning the locus folding by altering their SAW behaviour, include local crowding or additional internal contacts within these regions (such as presence of enhancers or local chromatin condensation). Taken together, our results suggest that the active/poised gene SBS polymer model, regardless of its over-simplicity, is sufficient to capture all the experimental measurements across the *HoxB* locus in mouse ES cells at the folding scale of tens of kilobases (~30-50kb, the sequence length of the FISH probes used). We do not exclude a role for CTCF occupancy in fine tuning the folding at shorter length scales. Our analyses hence support a scenario where distinct homotypic interactions between Active and Poised genes occur simultaneously across the vast majority of single cells in the heterogeneous ES cell population at distances lower than 100 nm; conversely, non-homotypic pairs of regions (i.e., all other eight cases) are seldom found below 200 nm.

**Figure 5.**
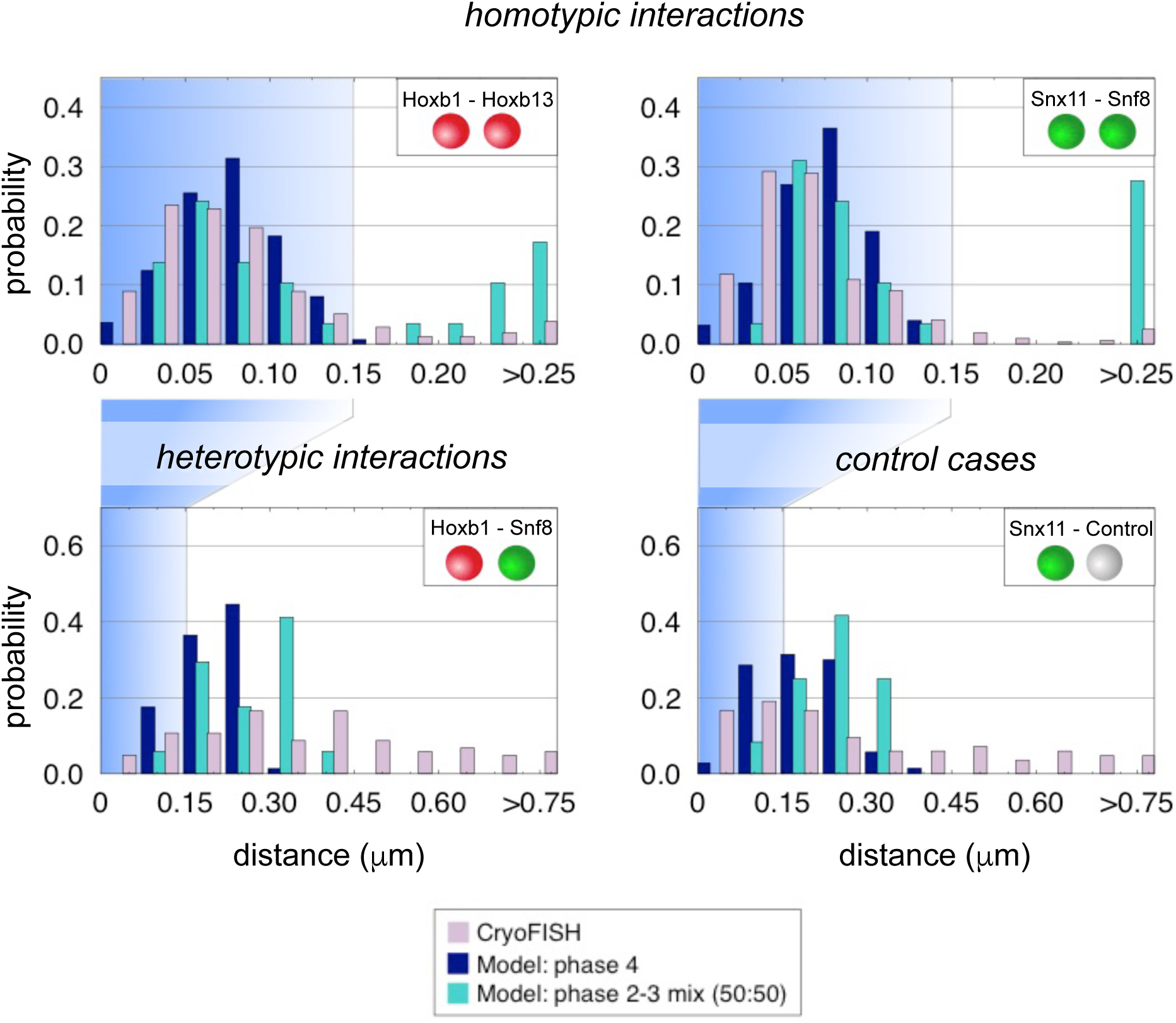
Agreement between cryoFISH imaging of the HoxB locus and the SBS homotypic interaction model extends to the distributions of inter-locus distances. **a)** Similar distance distributions were obtained experimentally from single cell cryoFISH and by polymer physics using an ensemble of SBS polymers with simultaneous (phase 4) homotypic interactions, although free fitting parameters were not used to optimise the match. Most (~90%) of the homotypic distances are lower than 150nm, in contrast with heterotypic and control probes distances that are much more broader (~70% above 150 nm). Note that the *x* axes scales are different between homotypic and the other cases. The profile of distances obtained experimentally is different from the profile obtained from an ensemble of SBS polymers produced from a mixture of polymers with only red-red and only green homotypic interactions (50:50 mixture of phase 2 and phase 3 polymers).

### Polymer conformations depend on the multiplicity of chromatin contacts

The mechanisms of folding identified within the *HoxB* locus suggest the formation of abundant contacts within each allele across the cell population, especially identifying long-range contacts between two distant regions separated by 730 kb, which contain active genes. To further test whether the same properties could be obtained by a more relaxed folding regimen, whereby each DNA binding region only loops with one other DNA binding region, we re-run the SBS model with the same red and green binder-binding site affinities but allowing each polymer bead to be bound at most by one binder (**Supplementary Fig. 8a**). Interestingly, we find that the contacts between the two Polycomb repressed regions containing *Hoxb1/Hoxb2* and *Hoxb13/AK078606* are still detected at a frequency of ~50% (in contrast with >75% of polymers when the multiplicity is maintained at six). Perhaps more strikingly, in these conditions of low multiplicity of contact, the two distant active regions, containing *Snf8* and *Snx11* respectively, separated by 730 kb (or 97 polymer beads), do not come together, a scenario disproved by experimental results. A correlation analyses between the locus proximities identified by cryoFISH and by the SBS modelling identifies the best correlation with phase 4 when binder multiplicity is 6 (**Supplementary Fig. 8b-g**). These results suggest that the folding of the *HoxB* locus is best explained by high abundance of local multiple contacts.

### Homotypic contacts between active and Polycomb repressed genes coincide with association with active transcription factories and Polycomb bodies

To further explore the nature of the homotypic contacts involving Active regions or Polycomb repressed regions, we tested whether the homotypic contacts observed coincide with the simultaneous association of Active or Polycomb repressed regions with nuclear domains containing the active form of RNAPII (also called transcription factories^2^) or containing Polycomb complex components (also called Polycomb bodies^3,6^). Our previous immuno-cryoFISH work had shown co-localisation of single active and Polycomb-repressed genes with RNAPII-S2p and Ezh2 in mouse ES cells^3^. Therefore, we combined cryoFISH of the active or poised genomic regions with immunodetection of the active (serine-2 phosphorylated) RNAPII or with Polycomb subunit Ezh2 (the catalytic subunit of PRC2; **Fig. 6**).

**Figure 6.**
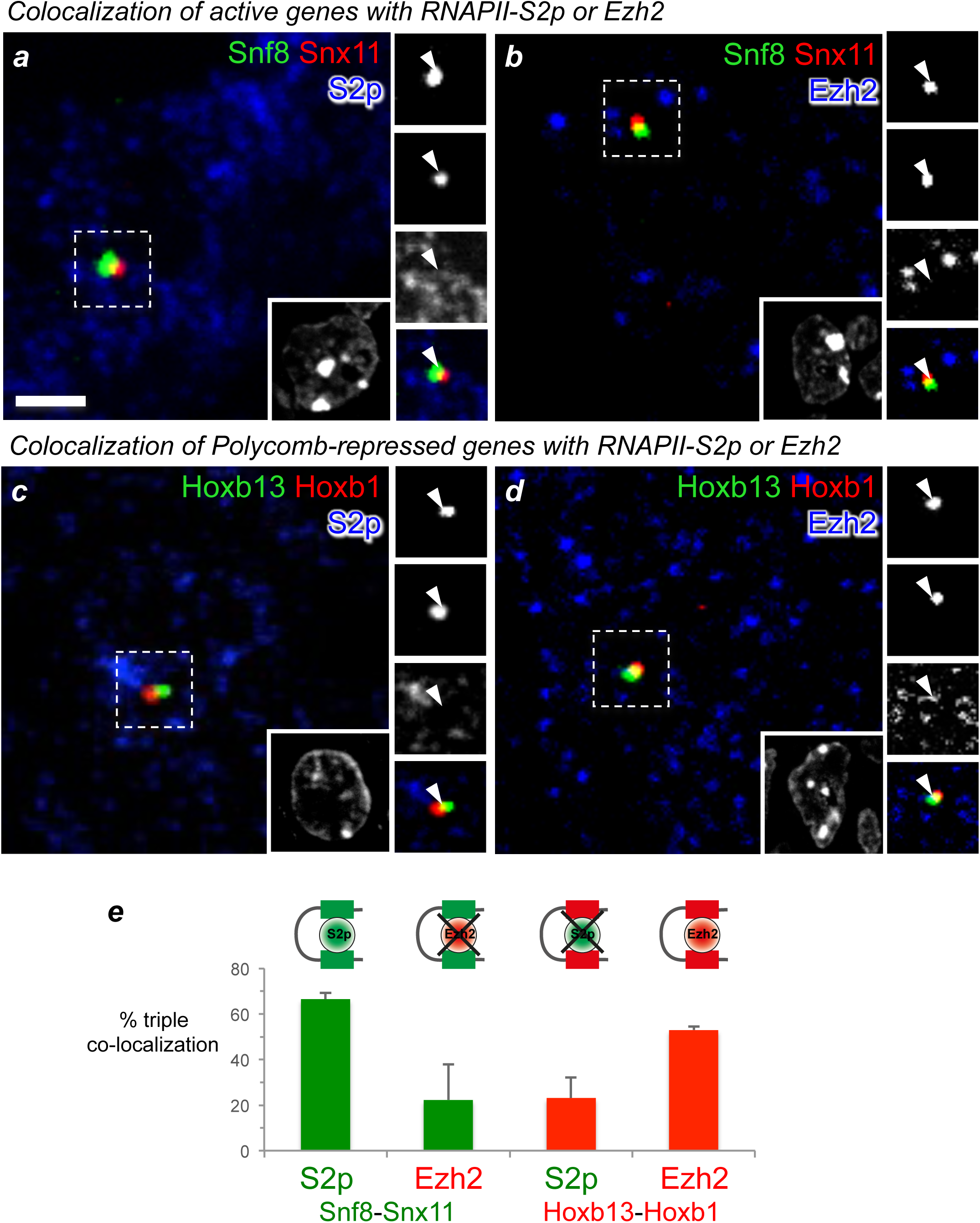
Homotypic interactions between active and poised genes coincide with their respective colocalisation with RNAPII-S2p and Polycomb complexes. **a-d)** The association of active and poised genes with active (S2p) RNAPII or Polycomb component Ezh2 was determined by immuno-cryoFISH. Before cryoFISH, cryosections (~150 nm thick) were indirectly immunolabelled with antibodies specific for RNAPII-S2p (**a**,**c**; pseudo-coloured blue) and catalytic Polycomb subunit Ezh2 (**b**,**d**; blue). The position of active genes (**a**,**b**: *Snf8 andSnx11,* pseudo-coloured green and red, respectively) or Polycomb-repressive genes (**c**,**d**: *Hoxb13* and *Hoxb1,* green and red) were imaged relative to RNAPII-S2p or Ezh2. Arrowheads indicate locus contact positions. Nucleic acids were counterstained with DAPI (blue). Bar, 2 μm. **e)** Frequency of association of *Snx8* and *Snf11* or *Hoxbl* and *Hoxb13* with active RNAPII-S2p and Polycomb enzyme Ezh2, markers of active transcription factories and Polycomb repressor complexes. Loci were scored as ‘‘co-localised’’ (fluorescence signals overlap by at least 1 pixel) or ‘‘separated’’ (signals do not overlap or are adjacent). Co-localisation of genomic regions containing active promoters *Snx8* and *Snf11* preferencially coincides with co-localisation with RNAPII-S2p. Conversely, colocalisation of Polycomb-repressed (poised) regions coincides with co-localisation with Ezh2, but not RNAPII-S2p.

We performed immuno-cryoFISH by triple labelling of RNAPII-S2p or Ezh2 with the two interacting Active (*Snf8/Ube2z* and *Snx11/Cbx1*; **Fig. 6a,b**) or Poised (*Hoxb1/Hoxb2* and *Hoxb13/AK078606;* **Fig. 6c,d**) regions. We find that when two Active regions colocalise with each other (*Snf8/Ube2z* and *Snx11/Cbx1*), they are also more likely to colocalise with RNAPII-S2p, but not with Ezh2 (**Fig. 6e**). In contrast, co-localised Polycomb-repressed regions (*Hoxb1/Hoxb2* and *Hoxb13/AK078606*) are more often found co-associated with Ezh2, but not RNAPII-S2p (**Fig. 6e**). This confirms that interactions among chromatin regions with similar epigenetic state (homotypic interactions between active or Polycomb-repressed genes) are favoured, with low-abundance of contacts of regions with different states (heterotypic interactions). Furthermore these results suggest that homotypic interactions between Active and Polycomb repressed promoters occur at transcription factories and Polycomb bodies, respectively.

## Discussion

Here, we investigated the contribution of gene promoter contacts to the 3D organisation of a 1Mb gene-dense region centred on the *HoxB* locus, by combining polymer physics simulations with imaging at the single-cell level. With the help of polymer modelling, we were able to identify critical genomic regions that could distinguish different mechanisms of locus folding. Single-cell cryoFISH results were consistent with formation of two kinds of homotypic contacts, dependent on promoter activation state alone, between active genes and between Polycomb repressed genes (**Fig. 7a**). We showed that the spatial proximity between active or between Polycomb-repressed genes coincide, respectively, with co-localisation with the active (Serine-2 phosphorylated) form of RNAPII or with the enzymatic component of PRC2, Ezh2. We also investigated other possible folding scenarions (**Fig. 7b**), in particular the hypothesis that CTCF alone could drive locus organization through formation of contacts involving CTCF binding sites^11-14^, but CTCF-driven folding alone was not compatible with FISH results, consistent with recent observations^10^.

**Figure 7.**
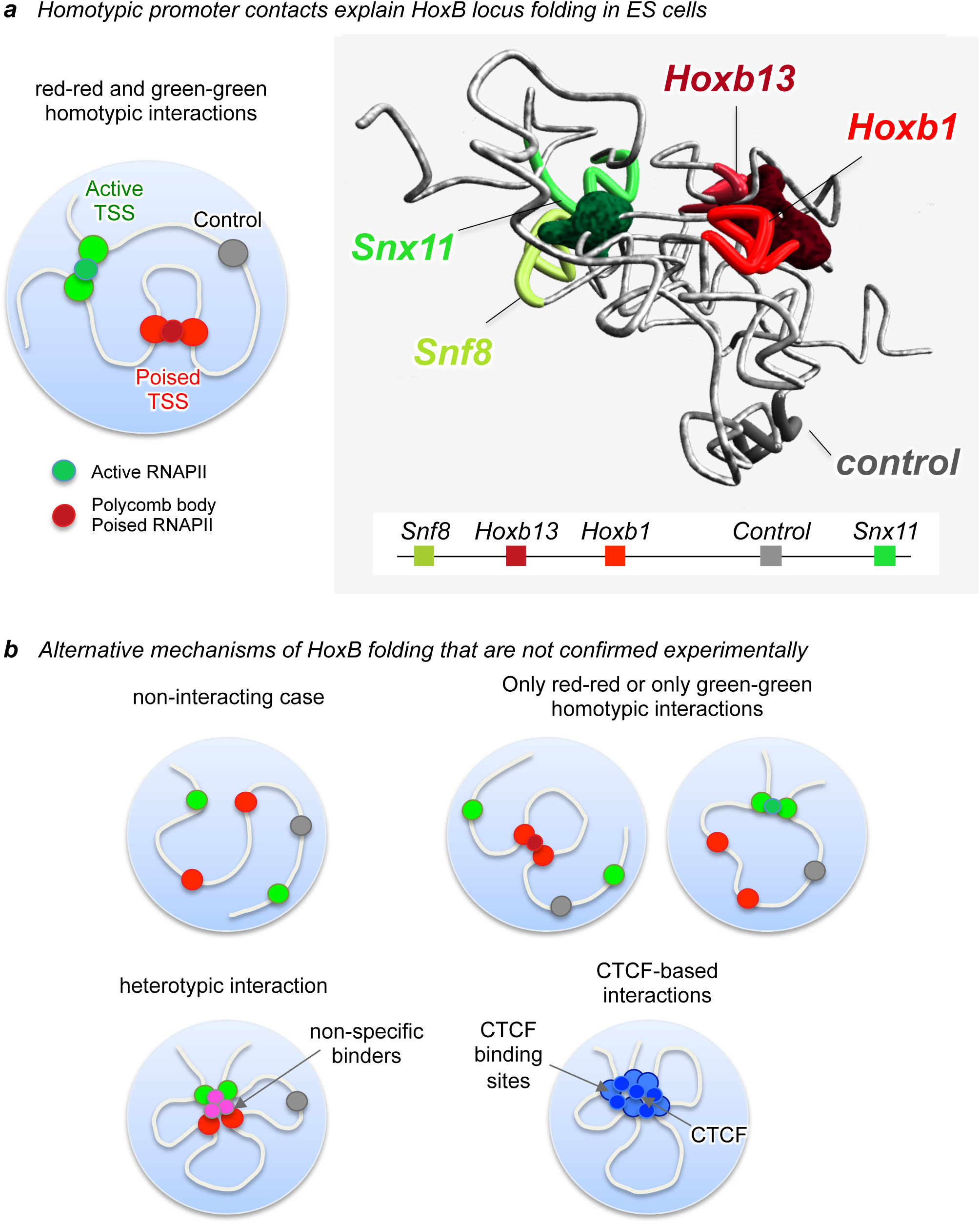
Summary model representing *HoxB* locus folding in mouse ES cells. **a)** The folding properties of the *HoxB* locus in mouse ES cells are explained, at the 30kb scale, by a polymer model based uniquely on homotypic interactions between active and poised gene promoters. Snapshot of the 3D conformation of the *HoxB* locus predicted by the phase 4 SBS model. Linear view of the *HoxB* locus regions tested by FISH. **b)** The alternative mechanisms of folding tested were not consistent with locus positioning measured in single cells by FISH: the non-interacting case where only random collisions, determined by relative genomic distance of chromatin sites, shape the locus architecture; two models where partial interaction among only one type of promoters is considered, i.e., where only active or only poised promoters have affinity with binders; a model based on heterotypic interactions between gene promoters, where each promoter has similar affinity with all binders (no active and poised promoters classification); and a model where the binding sites correspond to CTCF enrichment regions are the force which pulls together different parts of the locus

Our approach shows that 3D chromatin folding is driven by specific functional states across the chromosome fibre, and confirms that contiguous genomic regions are not necessary the closest in 3D space. This suggests that chromatin folding is more complex than a series of segregated local domains, with long-range interactions playing complex regulatory part via chromatin organization. More generally, the present study shows that the combination of *in-silico* polymer modelling, epigenetic feature analyses across the linear genome and high-resolution imaging of specific loci opens the way towards a deeper understanding of the relationship between chromatin architecture and chromatin states.

## Acknowledgements

We thank Kelly J. Morris for help growing the mouse ES cells, and Emily Brookes and Robert A. Beagrie for help. The work was supported by the Medical Research Council, UK (AP, SQX, IdS, MRB, DR), by the Helmholtz Foundation (AP, MB, ETT), and by the Berlin Institute of Health (AP, MB). This work was supported by grants to MN from CINECA ISCRA ID HP10CYFPS5. MN also acknowledges computer resources from INFN, CINECA, and *Scope* at the University of Naples.

## Author Contributions

A.P. and M.N. designed the project. M.B. performed the polymer modelling analysis.

S.Q.X. performed the wet-lab experiments and image analysis. I.d.S. and E.T.T. performed the bioinformatics analyses. D.R. provided conceptual advice and co-mentored the work of S.Q.X. with A.P.. M.R.B. developed the image analysis pipeline. A.P., M.N., M.B. and S.Q.X. wrote the paper, and all authors revised the manuscript.

## Online Methods

Methods, supplementary materials, statements of data availability and associated accession codes are available in the online version of the paper in the Supplementary Materials and Methods file.

